# Rare instances of non-random dropout with the monochrome multiplex qPCR assay for mitochondrial DNA copy number

**DOI:** 10.1101/2021.10.11.463983

**Authors:** Stephanie Y. Yang, Charles E. Newcomb, Stephanie L. Battle, Anthony YY. Hsieh, Hailey L. Chapman, Hélène C. F. Côté, Dan E. Arking

## Abstract

Mitochondrial DNA copy number (mtDNA-CN) is a proxy for mitochondrial function and has been of increasing interest to the mitochondrial research community. There are a number of ways to measure mtDNA-CN, ranging from qPCR to whole genome sequencing [1]. A recent article in the Journal of Molecular Diagnostics [2] described a novel method for measuring mtDNA-CN that is both inexpensive and reproducible. After adapting the assay for use in our lab, we have found it to be reproducible and well-correlated with mtDNA-CN derived from whole genome sequencing. However, certain individuals show poor concordance between the two measures, particularly individuals with qPCR mtDNA-CN measurements >3 standard deviations below the sample mean, which corresponds to roughly 1% of assayed individuals (Figure 1). After examining whole genome sequencing data, this seems to be due to specific polymorphisms within the D-loop primer region, at positions MT 338, 340, 452, 457, 458, 460, 461, 466, and 467. All individuals with a variant in at least one of these positions have non-concordant mtDNA-CN measurements. Meanwhile, variants observed at other positions within the primer region do not appear to cause dropout.

---

In particular, individuals from the U, L, and T mitochondrial haplogroups appeared to be more susceptible to failure. We classified discordant qPCR measurements as >3 standard deviations below the mean qPCR measurement (red dotted line), then used binomial logistic regressions to evaluate associations between discordant measurements and all haplogroups. Indeed, the U (p = 0.002), L1 (p = 0.041), L4 (p = 0.0003), and T (p = 0.015) haplogroups each had significantly greater individuals with discordant measurements. All other haplogroups, including L2 and L3, were not significantly associated with discordant measures. However, this could be due to limited power due to small sample sizes for certain haplogroups.

**Figure 1.**
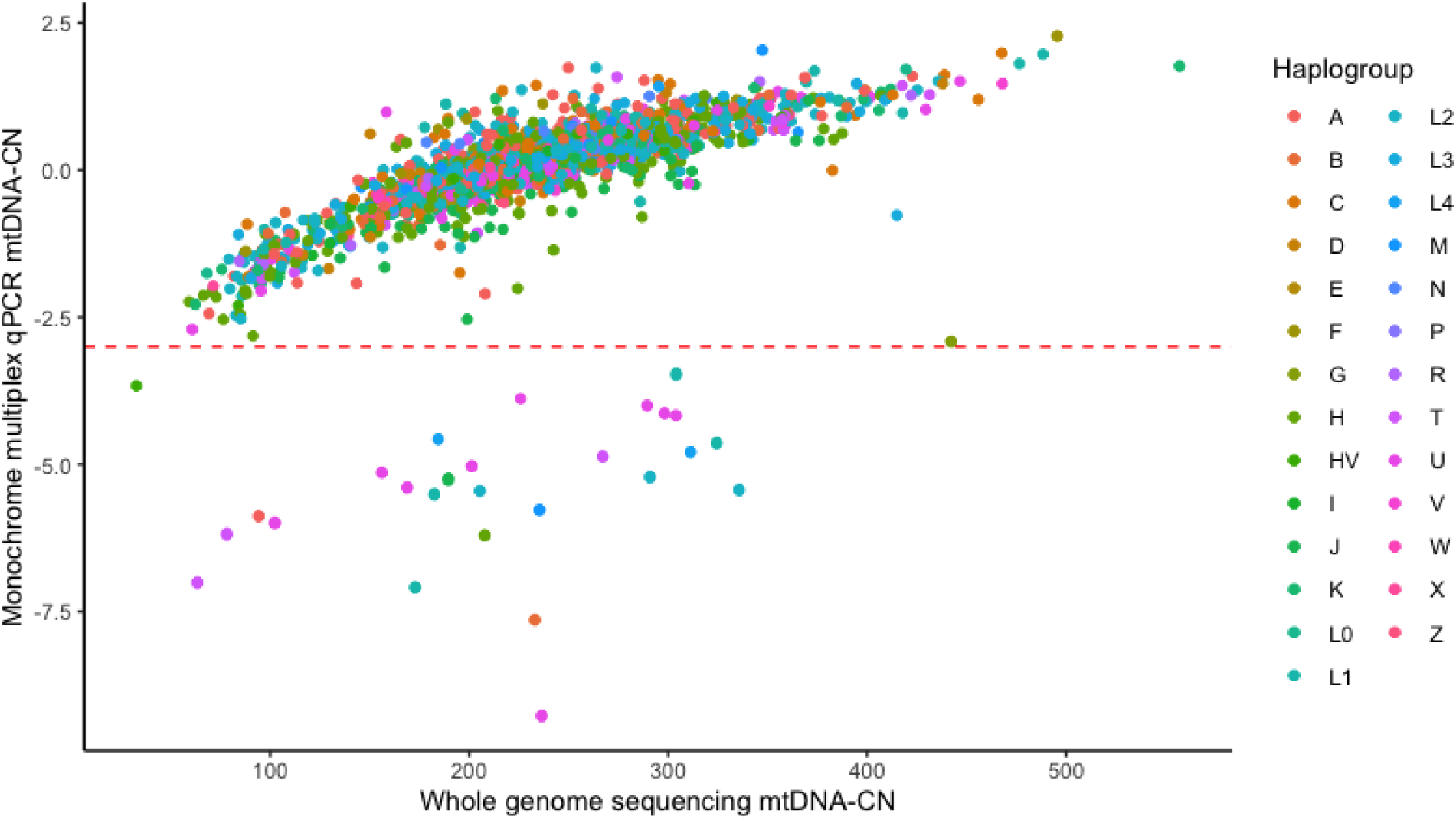
Discrepancy between the monochrome multiplex qPCR mtDNA-CN and the whole genome sequencing mtDNA-CN for 1,732 distinct individuals. Data are centered at 0 and scaled so that the standard deviation = 1. The dotted red line represents 3 standard deviations beneath the sample mean. Individuals in the U, L1, L4, and T haplogroups have a disproportionately higher risk of discordant measures between the two assays.

We hereby want to make other researchers aware of this non-random dropout, and of the need to confirm extremely low measurements obtained from this assay by using alternative assays that target other regions of the mitochondrial genome.

## Acknowledgements

This work utilized participants from the Multi-ethnic Study of Atherosclerosis (dbGaP accession: phs000209.v13.p3).

